# Parallel and scalable balanced minimum evolution algorithm for large-scale phylogenetic inference

**DOI:** 10.64898/2026.07.27.740982

**Authors:** Chris Tapo, Qiyun Zhu

## Abstract

The rapid growth of biological data demands scalable yet robust phylogenetic methods. Distance-based approaches are widely adopted not only in alignment-based molecular phylogenetics but also in emerging applications involving alternative distance measures. Balanced minimum evolution (BME) is an important principle in distance-based phylogenetic inference. A heuristic for BME using a taxon addition strategy was developed and provided by the FastME package, producing accurate tree topologies compared with alternative methods. However, beyond the original FastME implementation, there has been little development aimed at scaling this algorithm to the size of modern datasets. We present a significantly redesigned implementation of the BME algorithm that substantially improves computational efficiency while preserving mathematical equivalence. The key algorithmic improvement is the replacement of recursive tree traversal with flattened, incrementally growing arrays representing tree topology, traversal order and node properties. This design reduces traversal overhead, improves memory locality, and enables parallelization of the BME algorithm for the first time. Benchmarks show that our implementation delivers multiple dozen-fold speedup while consuming half the memory of FastME on datasets with tens of thousands of taxa. It can effectively analyze datasets of over 100,000 taxa, which are prohibitive with FastME. In addition, this implementation compares favorably in both efficiency and solution quality with a modern implementation of neighbor joining (NJ), an alternative BME heuristic using an agglomerative clustering strategy. Our BME algorithm has been released as part of the open-source Python library scikit-bio. It substantially broadens access to BME-based phylogenetic inference at scale.

## Introduction

The unprecedented accumulation of biological sequence data continues to push the computational limits of phylogenetic inference [1]. Distance-based methods remain an indispensable choice for analyzing ultra-large datasets [2]. They are applied not only as standalone tools for rapidly inferring clustering and evolutionary patterns, but also for the generation of starting trees to initiate and accelerate more complex phylogenetic optimization approaches [3]. Using pairwise distances as a proxy to aligned sequence characters enables phylogenetic analyses where alignments are not available, for example the MinHash distances among whole genomes [4], or distances in the embedding space inferred by protein language models [5], and also in emerging applications involving alternative distance measures, such as chemistry [6], linguistics [7], and communication [8].

Balanced minimum evolution (BME) is an important framework in distance-based phylogenetics. The minimum evolution principle seeks a tree that explains an observed distance matrix with the smallest total branch length. BME, introduced by Desper and Gascuel [9], calculates the tree length in a “balanced” way [10] such that sibling subtrees contribute equally, regardless of the size of each subtree (see Methods for details). Phylogenetic methods optimizing for the BME criterion produce accurate trees compared with alternative models [9], and according to some benchmarks even matching or exceeding the computationally intensive maximum likelihood and Bayesian methods [11, 12, 13]. The widely adopted neighbor joining (NJ) method is essentially a greedy heuristic for the BME criterion using an agglomerative clustering strategy [14, 15]. Another greedy heuristic, directly named as “BME”, was introduced by Desper and Gascuel [9] and implemented in the FastME package [15, 16]. This algorithm builds a tree by iteratively adding taxa to a growing tree. Specifically, each new taxon is inserted into a branch that gives the smallest balanced length for the updated tree (see Methods for details). The tree resulting from this progress can be further refined using tree rearrangement. This mathematically intuitive BME algorithm has a lower average-case time complexity than standard NJ. Initial tests on very small datasets (<100 taxa) showed that it produces slightly less accurate trees than NJ does [9], whereas later works on moderate datasets (500 taxa) revealed that its accuracy outperforms NJ [17]. Theoretical studies have shown that this BME algorithm is more robust than NJ to errors in the input distances and offers stronger guarantees of recovering the true tree [18, 19].

Despite these advantages, the application of BME at the scale of modern phylogenomics has been hindered by computational bottlenecks. FastME is capable of analyzing up to 2^16^ = 65, 536 taxa, a limit that is increasingly inadequate. Aside from this design decision, FastME’s algorithmic core was designed with a recursive tree-node data structure, which creates substantial overhead, impedes memory contiguity, and restricts the parallelization of tree traversal, collectively limiting the algorithm’s scalability. Later studies have primarily focused on advancing the mathematical understanding of the BME problem (BMEP), particularly in the context of integer linear programming [20] and polyhedral combinatorics [21], and methods for reducing the optimality gap under the BME criterion [22, 23, 24], instead of improving algorithmic efficiency and scalability beyond FastME’s baseline. Consequently, the applications of these later methods are limited to small datasets. While continuous efforts over decades have significantly accelerated NJ [25, 26, 27, 28], there is currently no BME implementation that effectively scales to datasets with hundreds of thousands of taxa.

We present a fundamentally redesigned implementation of the Desper and Gascuel [9] taxon-addition BME algorithm in the Python library scikit-bio [29]. Our approach replaces the recursive tree traversal with flattened, incrementally maintained arrays that permit direct indexing of nodes, clades, traversal order, and balanced average distances between subtrees. This representation improves memory locality, reduces redundant storage and arithmetic, and for the first time, enables effective parallelization of the BME algorithm. We demonstrate that this parallel BME algorithm achieves up to 82-fold speedup over the original in FastME on large datasets, while preserving mathematical equivalence. We further compare it with a modern implementation of NJ and show that the optimized BME algorithm is competitive in both efficiency and solution quality. These improvements make BME-based phylogenetics practical to substantially larger datasets and broaden access to scalable and robust tree construction within the Python scientific computing ecosystem.

## Results

### Algorithmic improvements

Our redesigned implementation of the Desper and Gascuel[9] taxon-addition BME algorithm is centered on an array-based representation of the growing tree. Instead of relying on recursive traversal of pointer-linked nodes, as the FastME code does, our method stores node relationships and properties in a series of flattened, preallocated arrays indexed by the order of insertion into the tree, such that new nodes are appended sequentially. Node indices as seen in a preorder traversal of the tree are stored in a separate array that is incrementally maintained during the iterative taxon-addition process. This preorder array converts recursive traversal into flat array iteration, and permits direct location of whole clades as contiguous blocks of nodes. This feature allows the algorithm to parallelize the dominant matrix update process by partitioning the tree into chunks of nodes and distributing them across worker threads. Our implementation can rapidly estimate the workload of individual nodes and whole clades, thereby generating approximately balanced chunks to improve parallelization efficiency. Additionally, while FastME stores both orientations of the symmetric balanced average distance between subtrees, our method stores only one orientation, as determined by preorder, and uses control flow to access this orientation without queries. The entire implementation has been carefully optimized to minimize redundant calculations. Our algorithm has a Python interface, with arrays managed by NumPy and compute-intensive loops written in Cython. Details of the mathematical process, data structures, algorithmic flow, and parallelization strategy are provided in the Methods section.

### Computational performance

We benchmarked our new parallel BME algorithm in scikit-bio against the original serial BME algorithm in FastME using random samples of prokaryotic 16S rRNA sequences from the SILVA database [30] (see Methods). Substantial gains in computational efficiency were observed (Fig. 1A, B, Table 1). While FastME’s tree-node-based C code remains more efficient for small datasets of 100 taxa or fewer, scikit-bio’s efficiency surpasses FastME at 200 taxa and above, and the gap widens as sample size increases. In serial mode, scikit-bio outperforms FastME from 7-fold at 1,000 taxa to 15-fold at 50,000 taxa. Parallelization further accelerates the algorithm, especially on large datasets. At 10,000 and 20,000 taxa, scikit-bio’s speedups over FastME are 45-fold and 66-fold, respectively. At 50,000 taxa, scikit-bio is 82 times faster than FastME, reducing the runtime from 3.0 hours to 2.2 minutes. While FastME is capped at 65,536 taxa, scikit-bio is able to analyze larger datasets. It analyzed 100,000 taxa in 10.6 minutes, and 200,000 taxa in 43.2 minutes.

**Figure 1.**
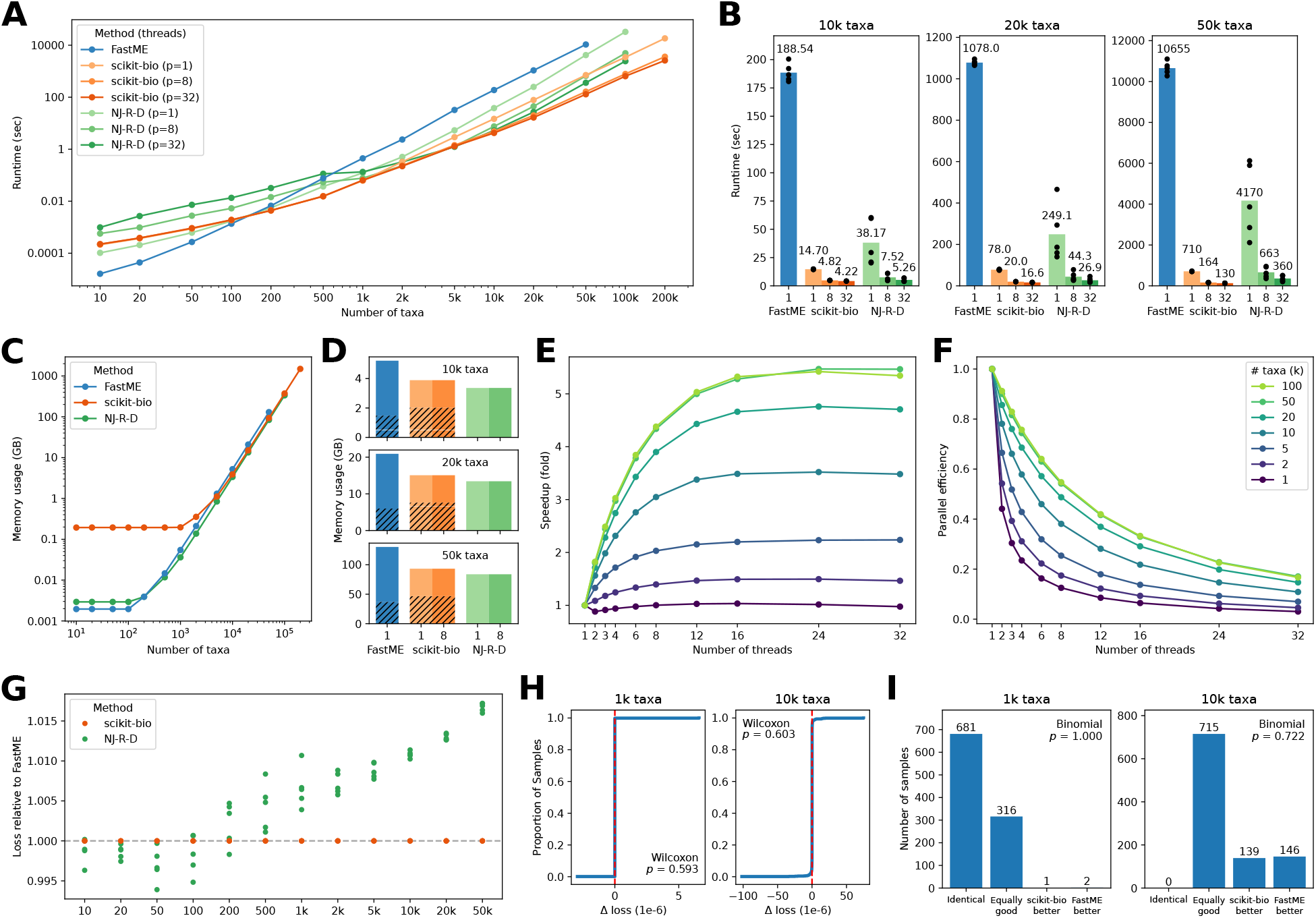
Performance of the parallel BME algorithm implemented in scikit-bio compared with the original implementation in FastME, and the neighbor joining algorithm (NJ-R-D) implemented in DecentTree. All three methods use double-precision floating-point arithmetic under the hood. **A.** Runtime of each algorithm over a series of taxon counts. Each data point represents the mean runtime of five random samples of taxa, each of which is calculated by the median of five runs, unless otherwise stated in Methods. FastME is serial only, whereas scikit-bio and NJ-R-D were run with one (serial), eight and 32 threads (*p*). “k” stands for thousand (same below). **B**. Comparison of runtimes of methods and thread counts at 10k, 20k and 50k taxa. Each data point represents the median of five runs per random sample. Each bar and the data label above it represent the mean of five samples. **C**. Memory consumption (maximum resident set size) of the three methods in serial mode. Refer to panel A for the meaning of data points. Values include both the memory usage of executing the algorithm and that of loading the distance matrix. **D**. Memory consumption of the three methods at 10k, 20k and 50k taxa, with one (serial) or eight threads. Refer to panel C for the meaning of the height of each bar. The hatched portion within each bar of FastME and scikit-bio represents the memory usage of loading the distance matrix. This value was not calculated for NJ-R-D. **E**. Speedup of scikit-bio in parallel vs. serial mode at a series of thread count with a series of sample sizes. Refer to panel A for the meaning of data points. **F**. Parallel efficiency (speedup vs. thread count) by thread count and sample sizes. **G**. Loss function (balanced tree length) of the output by scikit-bio and NJ-R-D relative to FastME (1.0, the dashed line). Values below 1 indicate better output than FastME. Each data point (ratio) was calculated using the same random sample of taxa. Five data points were calculated per sample size. **H**. Empirical cumulative distribution function (ECDF) of the difference (Δ) in loss of scikit-bio minus FastME for 1,000 random samples with 1k and 10k taxa. Two-sided Wilcoxon signed-rank test *p-*values are denoted. **I**. Numbers of samples out of 1,000 random samples in four categories: identical (FastME and scikit-bio output trees have the same topology), equally good (the two trees have different topologies but the same loss), scikit-bio better (its output tree has the smaller loss), FastME better (vice versa). Two-sided binomial test *p-*values are denoted.

**Table 1.** Computational efficiency of the parallel BME algorithm implemented in scikit-bio compared with the original implementation in FastME. Runtimes are in seconds. “vs. FastME” is the fold speedup (inverse of runtime ratio) over FastME, “Best” is the shortest runtime by any thread count. “Max. speedup” is the maximum speedup by any thread count over serial mode. “Max. threads” is the thread count achieving maximum speedup. See main text for the definition of “Practical threads”.

| Taxa | Serial runtime | Serial vs. FastME | Best runtime | Best vs. FastME | Max. speedup | Max. threads | Practical threads |
| --- | --- | --- | --- | --- | --- | --- | --- |
| 1000 | 0.062725 | 7.0922 | 0.060694 | 7.3295 | 1.0335 | 16 | 1 |
| 2000 | 0.32653 | 7.2152 | 0.21839 | 10.788 | 1.4952 | 24 | 12 |
| 5000 | 2.9103 | 11.113 | 1.3009 | 24.862 | 2.2372 | 32 | 12 |
| 10000 | 14.696 | 12.83 | 4.1746 | 45.164 | 3.5203 | 24 | 12 |
| 20000 | 78.045 | 13.813 | 16.399 | 65.737 | 4.7592 | 24 | 16 |
| 50000 | 709.87 | 15.01 | 129.79 | 82.096 | 5.4694 | 24 | 16 |
| 100000 | 3441.4 | – | 634.83 | – | 5.4209 | 24 | 16 |

Along with the remarkable gain of speed, scikit-bio’s memory consumption is moderately more efficient than FastME’s when analyzing large datasets (Fig. 1C, D). Because scikit-bio is a comprehensive Python library providing many features, it has a constant overhead of 160 MB of memory upon loading, which dominates the total memory consumption for analyses of 1,000 taxa or fewer. By comparison, the standalone C program FastME has an overhead of only 2.0 MB. However, scikit-bio uses less memory than FastME when analyzing 5,000 or more taxa, and this relationship converges to a constant factor. Specifically, scikit-bio uses 72% as much memory as FastME does at 20,000 or more taxa. If the memory cost of distance matrix loading is subtracted, this ratio drops to 50%. Additionally, there is no measurable difference in memory usage between serial and parallel modes of scikit-bio (Fig. 1D).

Further investigation of the parallelization behavior of scikit-bio reveals that its speedup is generally greater for larger datasets (Fig. 1E, F, Table 1). At 1,000 taxa, there is little speed gain with parallelization (1.03×). The practical thread count, defined as the minimum thread count achieving at least 95% of the highest speed with any thread count, is one, suggesting that multithreading is unnecessary at this dataset size. Parallelization begins to benefit at 2,000 taxa, achieving up to 1.50× speedup with 24 threads. The maximum speedup, 5.47×, was observed at 50,000 taxa and 24 threads. However, speedup does not further increase at 100,000 taxa (5.42×), suggesting a ceiling at this range of dataset sizes. Overall, the relative speed gain is largest at low thread counts and decreases as the thread count increases, with a plateau after 12 or 16 threads (Fig. 1E). Therefore, the practical thread count for datasets larger than 1,000 taxa is typically 12 or 16, although 24 or 32 threads can still provide modest additional gains (Table 1).

### Robustness of results

Next, we investigated the consistency between the tree outputs of scikit-bio and FastME. Although the two algorithms are mathematically equivalent, differences in design and implementation may lead to different trajectories of floating-point error accumulation and tie-breaking decisions, thus generating different outputs. We evaluated the output trees using balanced tree length—the loss function of the BME problem (smaller is better). Across sample sizes and random samples, scikit-bio achieves essentially the same loss as FastME does (ratio: 1.0 ± 1.5 × 10^−8^, mean and standard deviation, same below) (Fig. 1G). To examine this further, we conducted a more robust analysis by executing the two algorithms on 1,000 random samples of 1,000 or 10,000 taxa (Fig. 1H, I). Both methods produced nearly identical losses, with no statistical difference (two-sided Wilcoxon signed-rank test *p-*values = 0.593 and 0.603, respectively) (Fig. 1H). At 1,000 taxa, 68.1% of the pairs of scikit-bio- and FastME-generated trees had identical topology, while 31.6% of tree pairs had different topology but equal loss within eight decimal places (the default precision of FastME, same below), suggesting indistinguishable quality under the BME criterion. Only three of the 1,000 tree pairs had different losses, with scikit-bio yielding the smaller loss once and FastME twice, showing no statistical difference (two-sided binomial test *p-*value = 1.0, same below). At 10,000 taxa, no tree pair had identical topology, whereas 71.5% of tree pairs had equal loss. Among the remaining 285 cases, scikit-bio yielded the smaller loss in 48.8% of times and FastME in 51.2%, again showing no statistical difference (*p-*value = 0.722). When the losses were further rounded to five decimal places (the precision of the input distance matrices; see Methods), scikit-bio won in 56% and FastME in 44% of the 25 tree pairs with different topology (*p-*value = 0.690). These analyses suggest that the redesigned algorithm in scikit-bio achieves the same BME objective as FastME, with no detectable systematic bias in favor of either implementation.

### Comparison with neighbor joining

We additionally included neighbor joining (NJ) in the comparison, considering that NJ is an alternative heuristic to the BME problem, and it has been widely adopted and continuously improved [14, 15]. Specifically, we benchmarked “NJ-R-D”, an efficient re-implementation of the RapidNJ algorithm [27] in the DecentTree program [28]. Although other efficient NJ variants exist [31, 32], we chose this algorithm to represent NJ considering its adoption (DecentTree has been integrated into IQ-TREE for starting tree generation [33]), and its exactness (guaranteeing to build the same tree as standard NJ does). Our benchmarks show that scikit-bio’s taxon-addition BME algorithm is generally faster than NJ in both serial and parallel modes, and the advantage is more pronounced on larger datasets (Fig. 1A). However, NJ appears to scale better over threads than scikit-bio does. For example, at 50,000 taxa, scikit-bio’s BME is 5.9 times faster than NJ in serial mode, 4.0 times with eight threads, and 2.8 times with 32 threads (Fig. 1B). Meanwhile, NJ’s runtime varies greatly across datasets of the same size, in contrast to BME (either FastME or scikit-bio), which exhibits nearly constant runtime (Fig. 1B). On the other hand, the tested NJ implementation has slightly lower memory consumption than scikit-bio’s BME (90% at 50,000 taxa) (Fig. 1D), and a small overhead (3.0 MB) (Fig. 1C). With regard to BME loss, NJ is slightly better (0.18 ± 0.19% smaller loss) than taxon-addition BME on small datasets with 100 or fewer taxa, but it performs worse on medium-to-large datasets with 200 or more taxa, and the difference increases as the data size grows (Fig. 1G). For example, at 50,000 taxa, NJ’s loss is 1.7 ± 0.054% larger than BME.

### Software features

The BME algorithm is available through the `bme` function in the `skbio.tree` submodule. The input for this function is an instance of scikit-bio’s `DistanceMatrix` class, which provides a dedicated data structure and set of operations for handling pairwise distance data. scikit-bio supports parsing distance matrices in the classical Phylip format, plain TSV format, and a high-performance HDF5 format. In addition, multiple core scikit-bio functions produce distance matrices from various types of biological data. Examples include genetic distance calculation for aligned sequences, cophenetic distance calculation for taxa in a tree, and beta diversity calculation for community data. The output of the `bme` function is an instance of scikit-bio’s `TreeNode` class, which implements a recursive tree-node data structure. This class offers a large collection of methods for phylogenetic and general tree operations, including Newick format parsing, tree traversal, navigation between nodes, manipulation of tree topology, and calculation of tree and node metrics.

In addition to `bme`, the `skbio.tree` submodule offers the other three algorithms originally introduced by Desper and Gascuel [9] and implemented in FastME: GME, a taxon addition tree-building algorithm analogous to BME but using an unweighted ordinary least squares (OLS) framework; BNNI, an algorithm that refines an existing tree through nearest neighbor interchange (NNI) steps; and FastNNI, an OLS equivalent of BNNI. All four algorithms share the flattened tree array data structure and have been carefully optimized to improve computational efficiency. The `skbio.tree` submodule also provides an efficient implementation of the standard neighbor joining (NJ) algorithm, and a wrapper of SciPy’s `linkage` function for performing unweighted and weighted pair group method with arithmetic mean (UPGMA and WPGMA) tree building. Additionally, this submodule implements functions to calculate original and weighted Robinson-Foulds distances between trees. Together, these functionalities offer an integrated suite for large-scale, distance-based phylogenetic analysis.

## Discussion

We present a significantly redesigned and enhanced implementation of the Desper and Gascuel [9] taxon-addition balanced minimum evolution (BME) algorithm in the Python library scikit-bio [29] for distance-based phylogenetic reconstruction. Our implementation delivers substantial acceleration over the original implementation of this algorithm in the FastME program [16], while producing trees of equal quality under the BME criterion. This gain of performance is achieved through workload-balanced, race-free, and deterministic parallelization of the dominant matrix update process, optimized memory locality introduced by the flattened array structures, one-way storage of symmetric distances between subtrees, and reduction of repeated computation throughout the algorithm. These engineering improvements draw inspiration from past work that continuously advance computational phylogenetics (e.g., [28, 34]), and they contribute new inspirations for future advances, particularly in methods that share a distance matrix and/or iterative addition nature. The new BME implementation in scikit-bio empowers distance-based phylogenetic analysis through a modern Python interface. It not only produces trees of reasonable accuracy at scale, but also provides an efficient approximation to BMEP that will facilitate relevant mathematical research.

We further demonstrated that the parallel BME algorithm compares favorably in both efficiency and solution quality with the rapid neighbor joining (NJ) algorithm implemented in DecentTree [28]. With this comparison, we aim to highlight the value of BME as a competitive alternative to NJ for large-scale phylogenetic analysis. However, this comparison should be interpreted cautiously because it is not a comprehensive evaluation of all available NJ variants. Also, while it is meaningful to base tree quality evaluation on balanced tree length, since NJ is also a heuristic for BMEP, more insights will be obtained by incorporating other measurements such as topological distance to the ground truth or number of tree rearrangement steps needed to further improve the tree. Further comparison of BME with NJ and other methods will be interesting, especially on large datasets, and will be facilitated by our scalable implementation.

While the improvement of practical runtime is magnificent, the asymptotic complexity of the BME algorithm remains the same in our implementation, that is, average-case *O*(*n*^2^ log(*n*)) and worst-case *O*(*n*^3^) (see Methods). Future work may investigate potential heuristics to the exact taxon-addition BME algorithm, which itself is a heuristic to BMEP (analogous to various heuristics to NJ). A plausible direction has emerged from the current work: For subtrees with a sufficiently large topological distance (p) from the insertion point, 2^− *p*^ effectively becomes zero under IEEE floating-point arithmetic. The minimum *p* that satisfies 2^− *p*^ = 0 is 1075 for double precision, 150 for single precision, or 25 for half precision computing. Therefore, outside a constant radius of *p* − 1 around the insertion point, updates of balanced average distances between subtrees can be omitted (see Eq. (4)). That is, when the tree is sufficiently large, or more practically, when the desired numerical precision is limited, the time complexity of the BME algorithm effectively reduces to *O*(*n*^2^). The impact of numerical tolerance on final tree quality would need to be quantified.

Meanwhile, despite consuming half as much memory as FastME, our algorithm still requires *O*(*n*^2^) memory to store and process the balanced average distance matrix (see Methods), making it an important barrier to further scaling this method to even larger datasets, which are increasingly common in modern biological research. Overcoming this limitation should be a high priority for future work. Plausible technical solutions may include out-of-core computing (e.g., [34]) and sparse distance matrix adoption (e.g., [32]). Additionally, our parallel BME algorithm is bound by memory access patterns. While indexing the balanced average distance matrix using insertion order seems to be a necessary choice in order to avoid quadratic maintenance cost at each insertion (see Methods), the preorder scan of nodes by worker threads fragments the memory access of the matrix. Consequently, the parallel matrix update can become limited by cache locality and memory bandwidth, especially at high thread counts. Further improvements may require data layouts or scheduling strategies that better reconcile matrix storage and access. These considerations highlight a broader path forward for scalable distance-based phylogenetics, in which algorithmic and implementation designs must progress together.

## Methods

### Mathematical foundation of the BME algorithm

The balanced minimum evolution problem (BMEP) was introduced by Desper and Gascuel [9] based on the branch length estimation model developed by Pauplin [10]. It is a special case of the weighted least squares framework [35] that assumes the variance of distance estimate between two taxa is exponentially related to their topological distance in the tree [15]. BMEP seeks a tree topology *T* that minimizes the following objective function:

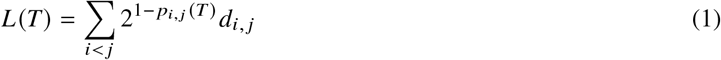

where *i* and *j* denote each pair of taxa. *d*_*i, j*_ is the observed distance between them provided in *D*, a symmetric distance matrix. p_*i, j*_ is the number of branches along the path connecting them in the tree *T* . *L* is referred to as the “balanced” tree length to be minimized under the minimum evolution principle.

Desper and Gascuel [9] proposed two complementary heuristics for the BME problem: 1. BME, a greedy algorithm that constructs a *de novo* tree through sequential taxon insertion. Later work referred to this algorithm as TaxAdd BalME (as in the FastME 2.0 program [16]), TaxAdd BME [24], Greedy BME [19, 23], or bGME [36]. We follow the original work to refer to this algorithm as “BME”, but want to remind readers to avoid the potential confusion between BME as an algorithm and BME as an optimality criterion. 2. BNNI, an algorithm that improves an existing tree through nearest neighbor interchange (NNI) moves under the BMEP criterion. FastME 2.0 [16] further implemented the subtree pruning and regrafting (SPR) method to complement NNI. This article focuses on optimizing the BME algorithm, although we also provide an optimized implementation of BNNI in the scikit-bio library.

The Desper and Gascuel [9] BME algorithm initializes with an unrooted 3-taxon tree *T*_3_, then iteratively inserts every subsequent taxon into the tree. Each taxon *k* is inserted into a branch in tree *T*_*k*−1_ that gives the smallest balanced tree length *L* (*T*_*k*_) after insertion (Fig. 2A). While computing *L* for all branches using Eq. (1) is expensive, Desper and Gascuel [9] showed that the change in *L* between two adjacent branches can be computed in constant time, thereby allowing all branches to be evaluated through a single tree traversal in linear time. Specifically, let *x* be an internal node joining three neighboring nodes *a, b* and *c*, moving the insertion point from branch {*x, a*} to branch {*x, b*} gives:

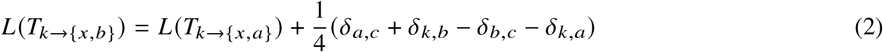

where *δ*_*i, j*_ denotes the balanced average distance between two disjoint subtrees rooted at nodes *i* and *j* . If both subtrees are leaves (i.e., they represent single taxa), there is *δ*_*a,b*_ = *d*_*a,b*_. Otherwise, *δ* is calculated by recursively averaging *δ*’s of sibling subtrees, such that they receive equal weight regardless of the numbers of taxa they contain:

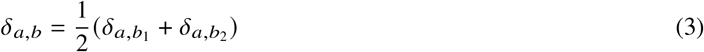

where *b*_*1*_ and *b*_*2*_ are the two sibling nodes branching from *b*. In each iteration, the *δ*’s between the new taxon *k* and all subtrees in tree *T*_*k*−1_ are calculated using the above equation through tree traversal, requiring linear time. The *δ* between any two subtrees is looked up from an incrementally maintained square matrix Δ, which stores the balanced average distances between all pairs of disjoint subtrees in *T*_*k*−1_ indexed by nodes. This matrix is updated during each iteration after the insertion of *k*. Specifically, only *δ*’s of subtree pairs into either of which *k* is inserted need to be updated. Consider nodes *x* and *y*, and *k* is inserted into the *y-*subtree, there is:

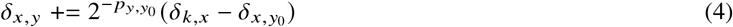

where *y*_*0*_ is the node on the further end of the branch into which *k* is inserted, relative to the root node *y*. 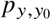 is the number of branches connecting *y* and *y*_*0*_ in the tree after insertion (Fig. 2A, in which 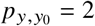).

**Figure 2.**
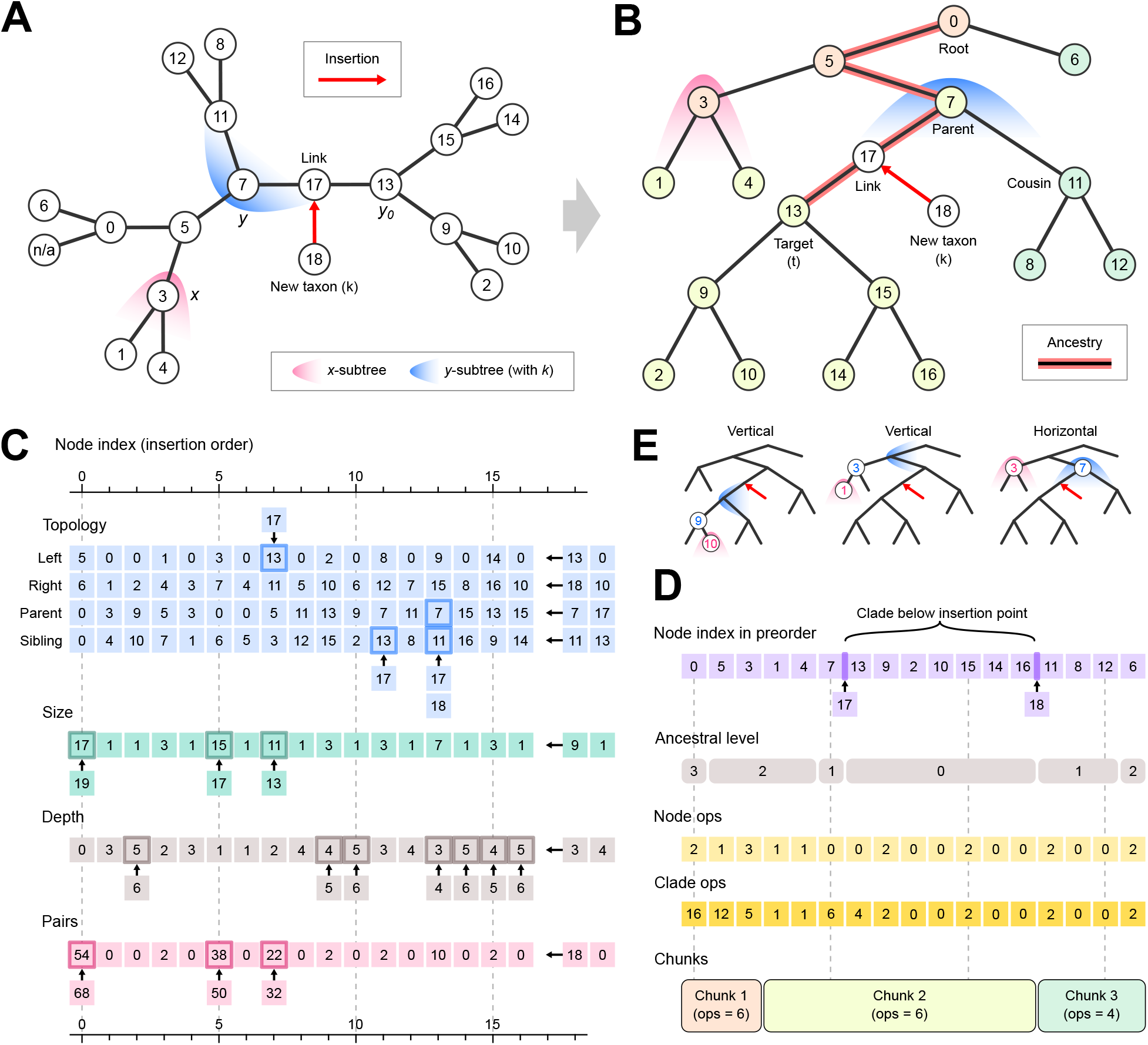
Illustration of the parallel BME algorithm using a dummy dataset of 11 taxa. The current scene depicts the insertion of the 11th taxon (*k*) into a tree that already has 10 taxa. **A.** Unrooted tree for mathematical representation. **B**. Rooted tree for computational representation. Lines represent branches and circles represent nodes, with labels denoting node indices in the order of insertion. The new taxon, with node index 18, is inserted into the branch connecting nodes 7 and 13, creating link node 17. Note that the first taxon, labeled “n/a” in the unrooted tree, is absorbed into the root node 0 of the rooted tree. Parabolas depict node-induced subtrees, with the focus indicating the root node and the opening indicating the branching direction. Node background colors indicate chunks used for parallelization. **C**. Flattened arrays representing nodes in insertion order. Numbers show the actual integer values stored in the arrays. The two cells appended at the right of each row represent the newly created link node and taxon node. Bordered cells with an arrow pointing from out-of-array cells represent values updated after taxon insertion. **D**. Flattened arrays representing nodes and node ranges in preorder. **E**. Examples of balanced average distances between subtrees that must be updated after taxon insertion. The node indices used in the Δ matrix are denoted. Note that in vertical pairs, the upper subtree is rooted at the parent of the indexing node.

The number of *δ* values updated at iteration *k* is *O*(*k* ·diam(*T*_*k*_)), where diam(*T*_*k*_) is the tree’s diameter, measured by the maximum number of branches between two leaves. The diameter is expected to have an average case of *O*(log(*k*)), for a tree that is balanced or follows the Yule–Harding model [37], which is commonly assumed in phylogenetics. The worst case is *O*(*k*), for a maximally skewed, comb-like tree. Collectively, the entire BME algorithm’s time complexity versus the number of taxa *n* is *O*(*n*^2^ log(*n*)) (average case) or *O*(*n*^3^) (worst case), whilst its space complexity is *O*(*n*^2^) due to the maintenance of the Δ matrix.

Our parallel BME algorithm implemented in scikit-bio follows the same mathematical process as the BME algorithm of Desper and Gascuel [9], but redesigns the data layout and update procedures to improve computational efficiency, as detailed below.

### Flattened arrays for efficient tree growth

Our BME algorithm is built around a set of flattened arrays representing tree topology and node properties (Fig. 2C). Each array element represents a node, indexed by the order of insertion into the tree. The first element always represents the root node. Although the growing phylogenetic tree is mathematically unrooted, rooting it for computation simplifies traversal and indexing. For algorithmic convenience, the leaf corresponding to the first taxon is omitted and absorbed into the root node, making the rooted tree strictly binary (Fig. 2A, B).

All arrays are preallocated with length 2*n*−3, because the (rooted) tree will have *n*−1 leaves and *n*−2 internal nodes. This allows memory to be reused throughout taxon addition and avoids repeated allocation and garbage collection. When a new taxon is inserted, two nodes are created and appended to the unused space of the arrays—a “link” node inserted into the branch connecting a target node and its parent, and a leaf node representing the taxon branching from the link node (Fig. 2A, B). The new taxon is always placed as the right sibling of the target. Tree topology is defined by storing node indices of four relationships of each node: left child, right child, parent and sibling. When the node is a leaf (i.e., a taxon), left child is set to 0 and right child is set to the taxon index. Upon insertion of a new taxon, only four elements around the insertion point need to be updated to represent the new relationships, requiring *O*(1) time (Fig. 2C).

Node indices sorted by a preorder traversal of the tree are stored in a separate array (Fig. 2D). This array converts tree traversal into array iteration, and allows each clade to be located as a contiguous block in preorder, starting from its root node and spanning its size (number of nodes), which is stored in a separate array (Fig. 2C). The preorder and size arrays are also maintained incrementally: At insertion, the link node is inserted in the preorder array immediately before the target clade, and the new taxon is inserted immediately after the target clade (Fig. 2D). This requires two memory-copy operations, which are *O*(*k*) but efficient in practice, and two assignments (*O*(1)). In the size array, 2 is added to every node on the ancestry of the target, from its parent to the root, because the link and taxon nodes become new descendants of each ancestor (Fig. 2B, C). This process is *O*(log(*k*)) on average. A separate postorder array is unnecessary. Although Desper and Gascuel [9] suggested using both preorder and postorder traversals for calculating *δ* values, we reason that the latter can be replaced by iterating the former in reverse order, which achieves equivalent bottom-up calculation because each internal node is visited after both of its children.

The algorithm also maintains a depth array, where each element *η* denotes the number of branches from the node to the root of the tree (Fig. 2C). This array facilitates quick calculation of the topological distance *p* between nodes without tree traversal. When a taxon is inserted, the depths of all nodes in the target clade increase by 1, requiring at most *O*(*k*) time. Additionally, the negative powers of 2 are precomputed up to *n* (largest possible topological distance +1), such that the factor 2^− *p*^ in Eq. (4) can be looked up without calculation.

The balanced average distance matrix Δ, which is the dominant term of the algorithm, is also preallocated with an order of 2*n* − 3 and indexed by the insertion order of nodes. This layout ensures that cells of each new link and taxon nodes are appended to the lower and right edges of the matrix (*O*(*k*) per iteration). Because balanced average distance is symmetric (*δ*_*a,b*_ = *δ*_*b,a*_), our algorithm only calculates and stores *δ*_*a,b*_ where *a* precedes *b* in preorder. This ordering is fixed for any pair of nodes since they are inserted, such that one never needs to calculate or store *δ*_*b,a*_ throughout the algorithm. Meanwhile, this ordering is directly known in the algorithmic flow without the need for explicit query or comparison. For example, an ancestor precedes its descendants, and a left sibling and its descendants precede a right sibling and its descendants. This design greatly improves memory efficiency for large datasets. One important note is that, under the rooted tree representation, the two subtrees involved in calculating *δ*_*a,b*_ are not always rooted at nodes *a* and *b*. Rather, one of them may be rooted at the corresponding node’s parent and branching upward. This notation will be clarified further below.

### The flat loops: finding the optimal insertion point

Each iteration in the outermost loop of the algorithm inserts the next taxon *k* into a branch that minimizes the resulting balanced tree length *L* (*T*_*k*_). It consists of the following steps. Our algorithm largely follows Desper and Gascuel [9], with modifications noted.

1. Calculate *δ*_*k,x*_, the balanced average distance between the new taxon *k* and every subtree in *T*_*k*−1_. In the rooted tree representation, each node *x* corresponds to two complementary subtrees that bipartition the tree at the branch connecting *x* and its parent: the lower subtree *x*^*L*^, which is the clade descending from *x*, and the upper subtree *x*^*U*^ = *T* \ *x*^*L*^, which is the subtree branching upward from *x*’s parent, with *x*’s sibling and grandparent as its two basal children (Fig. 2E). This notation follows Desper and Gascuel [9]. However, we would like to remind readers to avoid confusion, particularly in that subtree *x*^*U*^ is not rooted at node *x* but at its parent. Because the new taxon *k* may be inserted either below or above *x*, the algorithm calculates *δ*_*k,x*_ for both *x*^*L*^ and *x*^*U*^ for all nodes. The former is calculated bottom-up in reverse preorder (replacing postorder used in [9]), and the latter is then calculated top-down in preorder. Both use Eq. (3) and require *O*(*k*) time.
2. Determine the insertion point. The new taxon *k* will be inserted into the branch connecting a target node to its parent (Fig. 2B). The algorithm initializes the relative balanced tree length Δ*L* of the root node to 0, then iteratively calculates Δ*L* of the lower and upper subtrees of every non-root node in preorder following Eq. (2), using the just-computed *δ*_*k,x*_ values and the existing values stored in the Δ matrix. The node with the smallest Δ*L* is selected as the target node. Should there be ties, the first occurrence in preorder is selected. This procedure also requires *O*(*k*) time.
3. Update the Δ matrix. This is the dominant term, requiring *O*(*k* log(*k*)) time in the usual Yule–Harding average case and *O*(*k*^2^) time in the worst case. Its procedure will be described in the next subsection.
4. Update the flattened arrays to represent *T*_*k*_, using the procedures described in the previous subsection. They collectively require *O*(*k*) time.

Thus, steps 1, 2, and 4 are linear and are asymptotically cheaper than step 3. In practice, their main cost is memory access, especially after the parallelized step 3 has modified a large portion of the Δ matrix.

### The nested loops: updating the matrix

After inserting taxon *k*, the algorithm updates the Δ matrix, such that it reflects the balanced average distances between subtrees in the updated tree *T*_*k*_. Each cell value *δ*_*x,y*_ indexed by nodes *x* and *y* denotes the balanced average distance between the disjoint *x*-subtree and *y*-subtree. As discussed above, under the rooted tree representation, each node *x* corresponds to a lower subtree *x*^*L*^ and an upper subtree *x*^*U*^. However, given a pair of nodes, the choice of lower or upper subtree for either node is fixed by the tree topology, as exemplified below, and this relationship remains constant as the tree grows. Therefore, each *δ*_*x,y*_ value uniquely corresponds to one valid node/subtree combination. We assume the insertion point lies within the *y-*subtree, whereas the *x*-subtree remains unchanged (Fig. 2A). Following Eq. (4), the quantity added to *δ*_*x,y*_ is the product of a weight term, 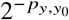 , and a difference term, 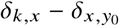. An intuitive implementation would compute both terms for each affected entry. Our algorithm reduces this cost by reusing intermediate quantities across nested loops.

First, consider the clade descending from the target node *t* (Fig. 2B). Within this clade, all ancestor (*y*)*–*descendant (*x*) pairs need to be updated, and *δ*_*x,y*_ represents the balanced average distance between *y*^*U*^ and *x*^*L*^. We refer to these as “vertical” pairs (Fig. 2E). They are updated by a nested loop in which the outer loop visits each node (*y*) in reverse preorder, and the inner loop visits the descendants *x* of *y* in preorder. For each outer-loop node *y*, the algorithm calculates the power 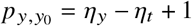, where *y*_*0*_ *is t’*s parent, and looks up the weight 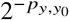 from the precomputed array. This value is calculated only once and then reused for all descendants of *y*. Similarly, the difference term of each node is calculated only once when the node is visited in the outer loop, and the value is reused later when the same node is visited as a descendant *x* in an inner loop. Thus, each vertical inner-loop calculation reduces to *δ* += weight × difference(*x*), requiring one multiplication, one addition, and one lookup.

Next, the algorithm traverses the ancestry of the target node *t*, from its parent toward the root. This traversal requires *O*(log *k*) time in the usual Yule–Harding average case. For each encountered ancestor *a*, the algorithm visits the cousin (i.e., the sibling of the previous ancestor), and all nodes descending from it (Fig. 2B). Each outer-loop node requires two inner loops: First, for an outer-loop node *y* and each of its descendants *x*, update *δ*_*x,y*_ concerning *y*^*U*^ and *x*^*L*^ (Fig. 2E). This resembles the vertical update inside the target clade, as described above, except that the power is now *p* = *n*_*y*_ − *n*_*t*_ + 2*v*, where *v* is the ancestry level, i.e., the number of ancestors encountered so far. This power, and hence the weight, is calculated once for each *y and reu*sed across its descendants *x*. Second, for an outer-loop *x* and each previous ancestor *y*, update *δ*_*x,y*_ concerning *y*^*L*^ *and x*^*L*^. We refer to these as “horizontal” pairs (Fig. 2E). The current ancestor itself is also subject to this update in the role of *x*, in which case *x*^*U*^ is concerned. For a previous ancestor at level *v*, the power is *p* = *v* + 1. The difference term, 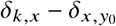, where *y*_*0*_ is *t*, is constant across previous ancestors for each *x*. Therefore, each horizontal inner-loop calculation reduces to *δ* += weight(*v*) × difference, again requiring one multiplication, one addition, and one lookup.

This organization preserves the mathematical update in Eq. (4), but avoids repeated computation of powers and difference terms.

### Parallelization of matrix update

To improve computational efficiency, our algorithm parallelizes the matrix update step by distributing outer-loop nodes across worker threads. Because each matrix entry to be updated is uniquely associated with one outer-loop node, the parallelization process is deterministic and free of race conditions. However, the workload associated with each outer-loop node varies substantially with tree shape. An even partition by node count could result in highly imbalanced workloads across threads. We use two strategies to mitigate this problem. First, OpenMP dynamic scheduling is used. Nodes are divided into chunks. The number of chunks is set to the number of available threads multiplied by an empirically chosen factor of 10. OpenMP assigns one chunk at a time to each thread and assigns the next available chunk when a thread completes its current work. Second, we estimate the workload of each outer-loop node and construct chunks with approximately balanced workloads.

As discussed above, the matrix update consists of vertical and horizontal inner loops, whose per-iteration costs are comparable. We consider each inner-loop iteration as one “operation”, or ops. For each node, the number of vertical inner loops equals the number of its descendants, or size − 1. The number of horizontal inner loops equals the number of previous ancestors, or *v* − 1, for nodes above the target, or 0 for nodes below the target. Therefore, the total number of operations per node is:

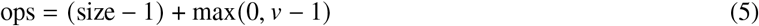

The size of each node is directly looked up from the size array. The level *v* is determined during the ancestry traversal. Because all nodes in a cousin clade share the same level, there is no need to keep a per-node level array. Instead, the algorithm divides the preorder array into segments representing contiguous blocks of nodes with the same level (Fig. 2D). Segment boundaries are placed at the target, each ancestor, and each right cousin (a left cousin immediately follows the ancestor with the same level). Given the size and level information, the algorithm can calculate the operations for each node while scanning nodes in preorder and assigning them to chunks (Fig. 2D).

We further reduce the cost of workload estimation and chunking by calculating the operations needed for entire clades. Because the preorder and size arrays locate each clade as a contiguous block, our algorithm scans nodes in preorder and “skips” over a whole clade if that clade’s workload fits into the remaining capacity of the current chunk. If the clade does not fit, the algorithm assigns the current node alone, either to the current chunk if capacity remains or to the next chunk otherwise. Each chunk is required to contain at least one node, even if that node’s workload exceeds the target chunk capacity. The scan then continues with the next node in preorder, which breaks the clade into smaller subclades. This procedure ranges from *O*(*k*) time in the worst case, when every node must be visited, to *O*(*c*) time in the best case, when each chunk receives one whole clade, where *c* is the number of chunks.

To support per-clade workload estimation, the algorithm maintains an additional array storing the total number of ancestor–descendant pairs within the clade descending from each node (Fig. 2C). This number depends on clade topology and is not a simple function of clade size. However, it can be updated incrementally after each insertion. As with the size array, only ancestors of the target node *t* need to be updated:

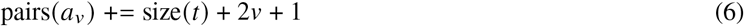

where *a*_*v*_ is the ancestor at level *v*. size(*t*) is before insertion. The constant 1 accounts for the link–taxon pair, and the factor 2 accounts for the ancestor–link and ancestor–taxon pairs at each ancestry level. Updating all ancestors requires *O*(log(*k*)) time on average and can be performed together with the size array update. This pairs array permits simple lookup of the vertical inner loop count of each clade. Meanwhile, the horizontal inner loop count is identical for all nodes in a clade. Therefore, the total number of operations per clade can be calculated as (Fig. 2D):

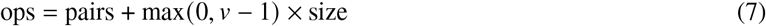

This estimate enables efficient workload-balanced chunking during a preorder scan (Fig. 2D). The description above focuses on the matrix update process, which is the dominant term of the entire BME algorithm. The actual implementation contains several additional engineering details. First, parallelization is performed in three phases: When the tree starts to grow, the algorithm runs serially to avoid parallelization overhead. When the tree grows to an intermediate size (default: 500 leaves), the nested matrix update process is parallelized. When the tree grows to an even larger size (default: 6,000 leaves), the flat computation for the optimal insertion point (see step 2 above) is also parallelized to further reduce runtime. It uses a simpler tree partitioning strategy and a deterministic tie-breaking strategy which are not detailed here. This three-phase execution plan ensures the algorithm’s efficiency for a wide range of data sizes. Second, the implementation includes special handling of the root node, the target node and its ancestors, and the created link and taxon nodes during matrix update to ensure that every *δ* value is calculated correctly under the current tree topology. These details are omitted from this article for conciseness, but they are documented in the source code through inline comments.

### Computer system for benchmarking

Benchmarking was performed on a Supermicro SYS-740A-T tower server configured with two Intel Xeon Platinum 8358 CPUs (32 cores and 64 threads per CPU, 48 MB cache per CPU, 2.60–3.40 GHz), 2 TB of DDR4 memory, and a 7.68 TB Intel D7-P5510 NVMe SSD. All tests were run on Linux Mint 22.1 (x86-64, kernel version 6.8.0-59).

### Benchmark dataset

SILVA release 138.2 [30] SSU Ref NR 99 sequences were used to benchmark the tree-building algorithms. This dataset consists of a precomputed multiple sequence alignment of 510,495 sequences by 50,000 sites. A series of numbers of sequences: 10, 20, 50, 100, 200, 500, 1000, 2000, 5000, 10000, 20000, 50000, 100000, and 200000, were randomly sampled from this dataset using seqtk. Five random samples per sample size were generated using seeds 0, 1, 2, 3, and 4, except for sample size = 200,000, for which only one sample was generated using seed 0 because of the extremely large size of the resulting distance matrix. For each sample, pairwise *p-*distances (fraction of differing sites) were calculated using scikit-bio’s `align_dists` function. We chose *p*-distance because this metric is bounded between 0 and 1, thus cross-comparable between samples, and it does not have the risk of producing undefined or infinite results, which could happen with more complex evolutionary distance metrics, considering the diversity of this dataset. Calculated *p*-distances were rounded to five decimal places to reduce disk usage. Sequence pairs with no shared sites were assigned the maximum distance *p* = 1.0. The same rounded distance matrices were used for all benchmarked programs.

### Benchmarking of FastME

FastME version 2.1.6.4 [16] (released on September 15, 2022) was downloaded from the official GitLab repository: https://gite.lirmm.fr/atgc/FastME/. We patched the source code of FastME to add two features related to bench-marking: 1. The program reports the precise time elapsed during the execution of the tree-building algorithm (C function `ComputeTree`), while excluding the time spent for loading the input distance matrix, writing the output tree, and other overheads. 2. It can optionally exit after loading the distance matrix and before building a tree. This permits measurement of memory usage needed for loading the distance matrix only, which can be subtracted from the memory usage of a complete analysis to obtain the extra memory consumed by the BME algorithm itself. The patched source code was compiled using GCC 13.3.0. We ran the FastME program using the following command: `fastme -T 4 -i input.phy -m B -w B -o output.nwk`, in which `-m B` represents the additive BME tree-building algorithm (“TaxAdd BalME”), and `-w B` represents branch length estimation under the BME criterion. Although FastME is capable of parallelization, this feature is only useful for calculating branch support statistics, whereas its BME algorithm cannot utilize parallelization. Therefore we set `-T 4` as a placeholder.

Each sample (taxon count by random seed; same below) up to 20,000 taxa was tested five times, and the median of the five runs is reported. When analyzing 50,000 taxa, each random sample was analyzed three times instead of five times, considering the extensive runtime. FastME cannot analyze more than 65,536 taxa, therefore it was omitted from the analysis of 100,000 and 200,000 taxa.

### Benchmarking of scikit-bio

The Python library scikit-bio was installed under Python 3.14.3 using the typical `pip install` command, which compiled the Cython code using GCC 13.3.0. To best represent the typical deployment of scikit-bio on an end user’s machine, we did not use any aggressive compiler flags (such as `-O3` and `-march=native`), although we acknowledge that compiler optimization may further accelerate the algorithm. The `bme` function of scikit-bio was executed to analyze an input distance matrix and generate an output tree. The elapsed time during this function call was reported, while excluding the time for loading the distance matrix into memory and writing the tree to a disk file. Note that the reported runtime also includes the time for converting the tree array structure into a tree-node structure at completion, which is nevertheless negligible compared with the tree-building process. Runs with distance matrix loading only but not tree building were executed to measure the memory usage of this process, which were then subtracted from the memory usage of a complete analysis to obtain the additional memory used by the BME algorithm. While scikit-bio’s BME algorithm provides native support for both single-precision (float32) and double-precision (float64) computing, we uniformly used double precision to ensure a fair comparison with FastME, which only supports double precision.

For the measurement of parallelization behavior in a multi-socket computer system, the benchmarking script binds threads to specific physical cores (`OMP_PLACES=cores`), binds worker threads to cores physically close to the parent thread (`OMP_PROC_BIND=close`), and pins the program to the cores and memory of the first NUMA node (`numactl -N 0 -m 0`). Each sample up to 100,000 taxa was tested using a series of thread counts: 1 (serial), 2, 3, 4, 6, 8, 12, 16, 24, 32. We tested up to 32 threads because that is the total number of physical cores per NUMA node. Five tests were run per sample per thread count, and the median of the five runs is reported. For the sample with 200,000 taxa, because the memory consumption exceeds the available physical memory of one NUMA node, we had to run the program without NUMA pinning. Considering the extensive runtime, we ran the test only once per thread count of 1, 8 and 32. Therefore, the 200,000-taxon tests were excluded from the analysis of parallelization behavior.

### Benchmarking of neighbor joining (NJ)

DecentTree version 1.0.0 [28] was installed from the Bioconda channel. The program is capable of reporting the elapsed time of the tree-building process, while excluding distance matrix loading. However, we were not able to modify the DecentTree source code to enable measurement of memory usage of distance matrix loading alone. Among the multiple NJ variants implemented in DecentTree, RapidNJ was shown to be highly efficient [28]. Instead of the default RapidNJ (a.k.a. “NJ-R”), which uses single precision computing, we chose “NJ-R-D”, its double-precision version to ensure a fair comparison with FastME and scikit-bio. The parallelization settings in the benchmarking script were identical to that of scikit-bio. We ran the DecentTree program using the following command: `decenttree -nt` $`p -t NJ-R-D -in input.phy -out output.nwk`, in which `$p` is the number of threads. Each sample up to 50,000 taxa was tested using 1, 8 and 32 threads, and the median of five runs per sample per thread count is reported. At 100,000 taxa, only one run per sample per thread count instead of five was performed, considering the extensive runtime. Tests of 200,000 taxa were omitted for the same reason.

## Code and data availability

The parallel BME algorithm has been incorporated into the scikit-bio package, available at: https://scikit.bio. Specifically, this algorithm is available from the `bme` function of the `skbio.tree` submodule. The source code of scikit-bio is licensed under BSD-3 and hosted at the public GitHub repository: https://github.com/scikit-bio/scikit-bio. Version 0.7.3 of scikit-bio, which started to ship the algorithm presented in this manuscript, has been published at the Python Package Index (PyPI): https://pypi.org/project/scikit-bio/0.7.3/. The benchmark scripts are available at: https://github.com/qiyunzhu/skbio-bme-bench.

## Author contributions

C.T. conceived the study, implemented a prototype of the BME algorithm, and instructed Q.Z. on the mathematical foundation of BME. Q.Z. redesigned and implemented the production code of the BME algorithm, performed benchmarking, and drafted the manuscript. Both authors contributed to designing, coding, and writing.

## Acknowledgments

We thank Dr. Siavash Mirarab for valuable comments on the scope of this study. We thank Matthew Aton for reviewing the code and performing continuous integration of the software package. This research was supported by the U.S. Department of Energy, Office of Science under award DE-SC0024320. This work utilized the computational resources of the Sol Supercomputer at Arizona State University.

## AI disclosure

Various AI models were used during the study as chatbots to search the literature, discuss algorithmic and experimental design, suggest code examples, and review code and the manuscript. However, AI models were not used as coding agents to directly generate or test code. The algorithm presented in this work was primarily human-written.

